# Getting stuck in a rut as an emergent feature of a dynamic decision-making system

**DOI:** 10.1101/2020.06.02.127860

**Authors:** Matthew Warburton, Jack Brookes, Mohamed Hasan, Matteo Leonetti, Mehmet Dogar, He Wang, Anthony G. Cohn, Faisal Mushtaq, Mark A. Mon-Williams

## Abstract

Human sensorimotor decision-making has a tendency to get ‘stuck in a rut’, being biased towards selecting a previously implemented action structure (‘hysteresis’). Existing explanations cannot provide a principled account of when hysteresis will occur. We propose that hysteresis is an emergent property of a dynamical system learning from the consequences of its actions. To examine this, 152 participants moved a cursor to a target on a tablet device whilst avoiding an obstacle. Hysteresis was observed when the obstacle moved sequentially across the screen between trials, but not with random obstacle placement. Two further experiments (n = 20) showed an attenuation when time and resource constraints were eased. We created a simple computational model capturing dynamic probabilistic estimate updating that showed the same patterns of results. This provides the first computational demonstration of how sensorimotor decision-making can get ‘stuck in a rut’ through the dynamic updating of its probability estimates.

**Significance Statement:** Humans show a bias to select the organisational structure of a recently carried out action, even when an alternative option is available with lower costs. This ‘hysteresis’ is said to be more efficient than creating a new plan and it has been interpreted as a ‘design feature’ within decision-making systems. We suggest such teleological arguments are redundant, with hysteresis being a naturally emergent property of a dynamic control system that evolved to operate effectively in an uncertain and partially observable world. Empirical experimentation and simulations from a ‘first principle’ computational model of decision-making were consistent with our hypothesis. The identification of such a mechanism can inform robotics research, suggesting how robotic agents can show human-like flexibility in complex dynamic environments.

## Introduction

Humans are creatures of habit and often repeat behaviours - despite the selected action having a higher cost than an available alternative. This propensity can be seen when humans continue to use the road well-travelled when moving between two buildings even after construction work has created a shorter route. The phenomenon is particularly remarkable because adult humans are generally so adept at selecting optimal movement patterns (Trommershäuser et al., 2008). Indeed, the ability of humans to rapidly and efficiently execute actions far exceeds the capabilities of even the most sophisticated robotic systems (Dogar & Srinivasa, 2012). The incredible repertoire of skilled behaviour in humans reflects the presence of learning processes that have been trained over the countless occasions when adults have interacted with the external world. These myriad interactions allow the human nervous system to accurately estimate the costs associated with various behaviours and thereby select an optimal (or close to optimal) action when presented with a goal directed task. The issue of relevance within this manuscript relates to the observation that adult humans will select different options on different occasions as a function of whether the choice is made *de novo* or following a previous successful action – despite the choices having the same relative costs on both occasions.

The tendency to show a bias towards a previously selected action plan can be described as ‘hysteresis’ (or the sequential effect) and is well-studied. Hysteresis effects have been found in grasp selection (Cohen & Rosenbaum, 2004, 2011; Dixon et al., 2012; Kelso et al., 1994; Kent et al., 2009; Rosenbaum & Jorgensen, 1992; Schütz et al., 2011; Short & Cauraugh, 1997; Weigelt et al., 2009), hand selection (Rostoft et al., 2002; Schweighofer et al., 2015; Weiss & Wark, 2009), and hand path priming experiments (Jax & Rosenbaum, 2007, 2009; van der Wel et al., 2007). However, there are no satisfactory explanations to account for this phenomenon. In fact, most explanations are teleological in nature: it is proposed that modifying a previously used action is more cognitively efficient than planning from scratch, so hysteresis exists to increase planning efficiency (Meulenbroek et al., 1993; Rosenbaum et al., 2007; Schütz & Schack, 2019; Weiss & Wark, 2009), as indexed by reduced reaction times (RTs) when using the same action as previously (Valyear et al., 2018). The problem is that such interpretations do not provide a principled account that can explain when hysteresis will occur, why its magnitude differs under different task constraints, or why its presence is a function of the costs of the available choices.

We propose that the process of ‘getting stuck in a rut’ is an emergent property of a decision-making system that dynamically learns from the consequences of its actions. In order to deal with unpredicted changes in the world (and adapt to novel environmental states), an efficient system must frequently update its estimates of the success probabilities associated with a given action (in Bayesian terms, the system must continually update its priors). To update these estimates, humans must use feedback about the outcomes of their actions – actions which cause the environment to transition to a new state. We suggest that this principle – the updating of success probabilities – will naturally result in a system that shows hysteresis. In fact, it is common practice in computer science to model the environment as a POMDP (POMDP; Kaelbling et al., 1998) when designing agents that need to act under uncertainty. In a POMDP, the agent does not directly observe the environment’s state but receives an observation which is a function of the state of the environment following an action executed by the agent. POMDPs reflect well the challenges faced by the human nervous system which must infer the hidden states of the environment from the sensory inputs that follow an action (as a Markov blanket separates the nervous system from the external world; Friston, 2010). The important point from the perspective of this manuscript is that the external world is not static and this means that a human agent must frequently update its internal representation (i.e. the approximate conditional density on the causes of sensory input) in order to act optimally in a noisy and changing world. The dynamical updating of the internal representation enables the human to predict the sensory input that will result from a generated action. The ability to make accurate predictions allows a human to generate an action that will produce a desired change in the sensory input (i.e. achieve a goal-directed change in the external environment). It follows that efficient action selection requires frequent updating of the success probabilities associated with a given action. We hypothesised that this updating would produce hysteresis as an emergent property of the dynamical learning system.

In order to explore hysteresis, we needed to design a canonical task that would allow us to parametrically vary critical task parameters, reflect a naturalistic action, and produce data amenable to computational modelling. We also needed a task that would allow us to examine behaviour on a trial-by-trial basis so that we could address issues relating to the frequency of updating (e.g. whether we would observe hysteresis on a trial-by-trial basis or whether it was only manifest after a number of iterations of a given action structure). We decided to use aiming movements (moving an end effector from a start point to a target location) to meet our task requirements. We therefore created a simple multi-trial sensorimotor decision-making task in which participants needed to move around an obstacle (left or right) to hit a target (where the reward was the same for each choice). We manipulated obstacle position on each trial across blocks such that it either moved systematically across the screen or was randomly positioned across trials (Figure 1). We predicted that participants would show a strong bias towards repeating previously selected actions in the sequential condition, even when the obstacle position indicated an alternative route would be preferable. We expected that this effect would be diminished in blocks where obstacle position moved randomly across the screen.

**Figure 1.**
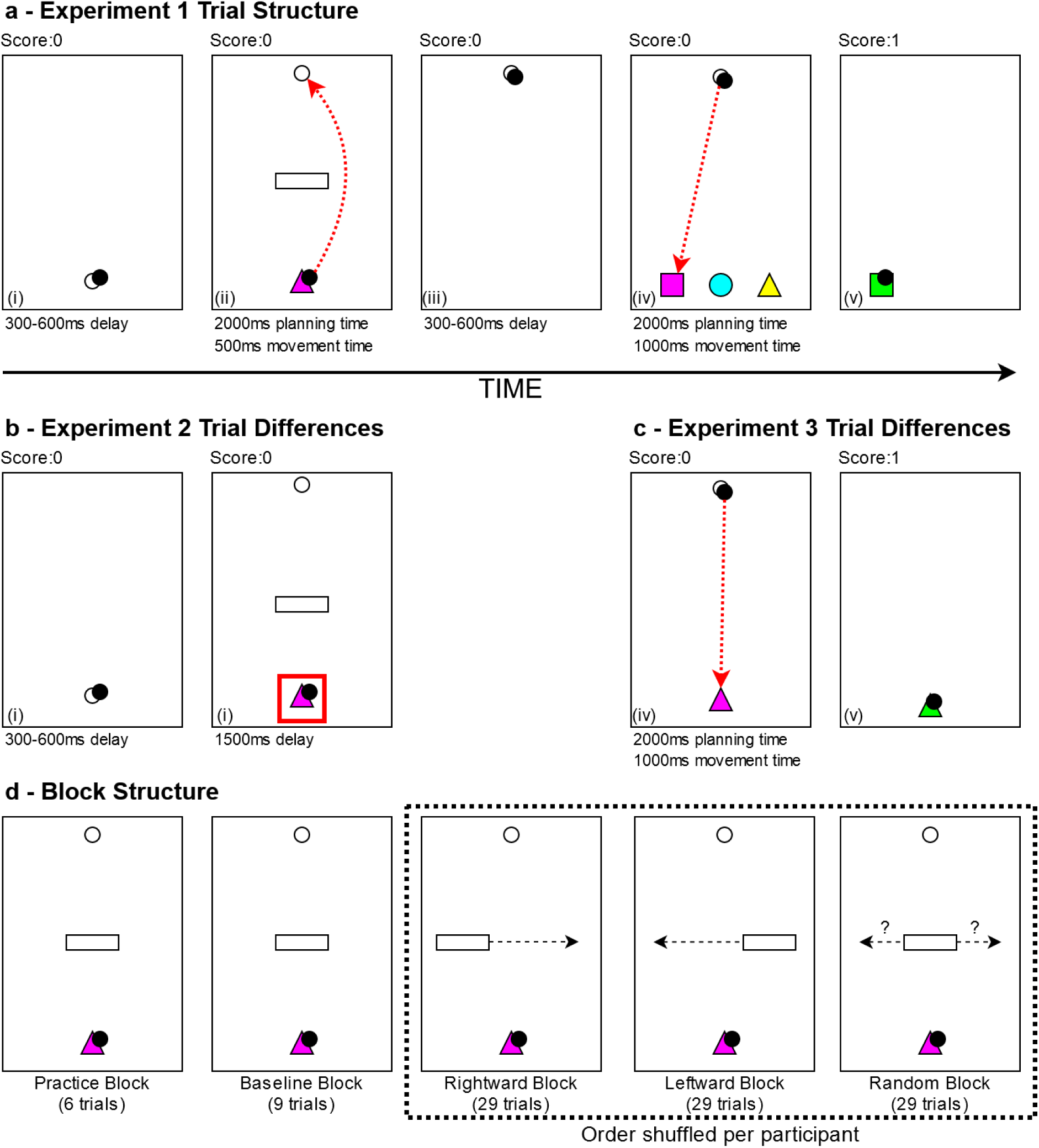
Trial and block structure of the experiments. (a) A complete trial for Experiment 1. Red dashed lines indicate potential movement trajectories and filled circle indicates movement endpoint. In Step (i), participants moved to a start-point and waited 300-600ms until it changed to a colour and a shape indicating the target shape colour. Simultaneously, an obstacle and a checkpoint appeared. Participants were allowed 2000ms planning time at the start-point before moving around the obstacle to the checkpoint (<500ms) and waited 300-600ms until 3 targets appeared. Participants were allowed up to 2000ms in the checkpoint before moving to the target that matched the colour shown at the start-point (<1000ms). (b) For Experiment 2, step (i) of Experiment 1 was replaced by two steps. Participants moved to a start-point and were immediately shown a red box around the start-point, indicating they could not leave. After a random 300-600ms delay, the stimuli were revealed but the red box remained on screen for a further 1500ms. (c) For Experiment 3, steps (iv) and (v) of experiment 1 were changed so only one target was revealed, of the same colour and shape as the start-point. (d) The block structure of the experiments. Participants completed a practice and baselining block, where the obstacle was always central to the screen, before completing a shuffled order of the Rightwards block (obstacle moves from the left to the right between trials), Leftwards block (the obstacle moves from the right to the left between trials), and Random block (obstacle moves randomly between trials).

We wished to explore whether the empirical data generated through our empirical investigations could be captured by a model that incorporated dynamic probabilistic estimate updating (i.e. whether hysteresis could be captured through a POMDP type model). We therefore created a model of human decision-making (Figure 2) that included a trial-by-trial update of the success probabilities associated with one action versus another. The goal of the model was to simulate how an agent would respond to a choice between two options that both allow a given goal to be achieved but have different costs. The output of the model was an action that would cause the environment to transition (with a given probability) to a new state. The model was arranged such that after an action is executed the agent receives an observation which is a function of the new environmental state. This input was then used to update the success probabilities associated with the action. In many choice tasks, there is also a difference in the reward associated with the options (Dreher & Tremblay, 2009; Gold & Shadlen, 2007; Mushtaq et al., 2016), so our model incorporated an estimate of the reward to allow future studies to explore behaviour in such tasks (but in the reported experiments the reward was identical across options).

**Figure 2.**
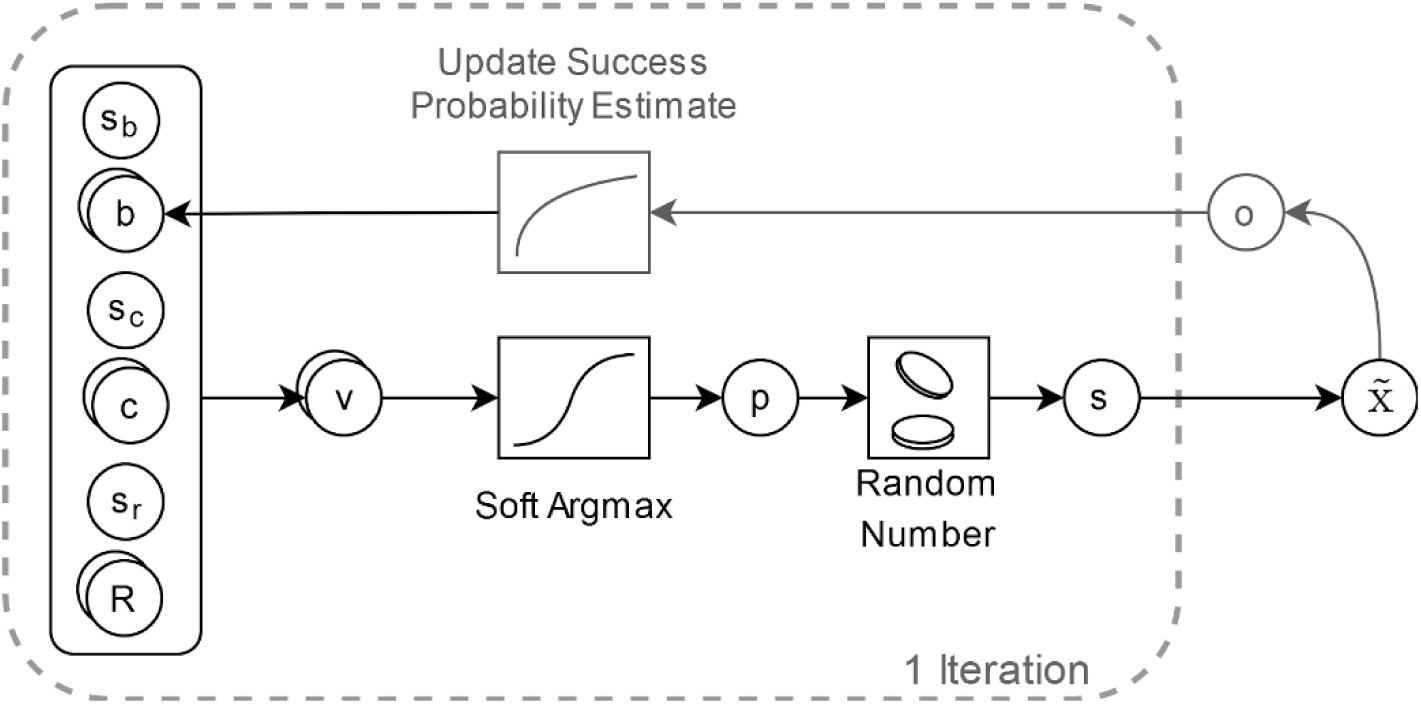
A probabilistic choice model for action selection. In a trial, the ‘value’ for the two actions (going left or right around the obstacle) is calculated from the current costs, rewards and biases built up over previous trials. The values are input to the soft argmax function which gives the probability of selecting the left action. A random number is uniformly sampled and if it is below the probability of the left action then left is selected, otherwise right is selected. The outcome of executing the associated action is observed and the selection and outcome are used to update the biases for each action according to a reinforcement learning rule.

We were also interested in exploring the dynamical aspects of decision-making under temporal constraint. Converging evidence suggests that the decision-making process is governed by neural circuits that accumulate noisy evidence for possible options over time, with a decision triggered when sufficient evidence is accumulated to cross an action threshold (Bogacz et al., 2006; Brody & Hanks, 2016; Gold & Shadlen, 2007). It seems reasonable to suppose that the action threshold will be a function of the available time period within which a decision must be made (i.e. temporal constraints will push the system towards making a choice that might be different were more time available to weigh up the respective costs of the different options). Notably, the existence of evidence accumulation processes predicts that actions will be selected more rapidly (i.e. RTs will decrease) when there is a bias towards one action versus another. This suggests that RTs will be faster in the presence of hysteresis. Once more, it is important to emphasise that we are proposing that hysteresis is an emergent property of a dynamic learning system where faster RTs are a useful by-product of the system’s organisation rather than the planned product of a system designed with an inbuilt function to produce hysteresis.

On the assumption that evidence accumulation processes are a core component of sensorimotor decision-making, we anticipated finding RT differences as a function of the magnitude of hysteresis associated with a given task. We further hypothesised that relaxing the temporal constraints of the task would attenuate the size of the hysteresis effect (as the available time can be used to more fully evaluate the costs of either action, reducing the reliance of the decision-making system on previous successes and failures to inform current action selection). In Experiment 2, we directly manipulated the temporal constraints of the task by creating a ‘waiting period’ before which an action could be executed. In Experiment 3, we indirectly manipulated the temporal constraints by decreasing the ‘higher order’ cognitive demands of the task. We reasoned that decreasing the cognitive constraints would allow the task goal to be identified more rapidly and thereby create a longer period in which the respective costs of the alternative actions could be computed.

## Methods

### Participants

In Experiment 1, 152 adults (41 males, 100 females; mean age 22.51 years, range 18– 39 years; 139 self-reported right-handed; eleven participants did not report age or gender) were recruited as part of a larger motor control project. Participants for Experiment 2 (n = 20, 1 male, 19 females; mean age 19.09, range 18-20 years; 20 self-reported right-handed) and Experiment 3 (n = 20, 1 male, 19 females; mean age 18.86 years, range 18-20 years; 18 self-reported right-handed) were recruited through word of mouth from the University of Leeds undergraduate population. All had normal or corrected-to-normal vision and provided informed consent to participate. Participation in these studies was incentivised through remuneration of £2 on completion of the experiment. Ethical approval was obtained from the University of Leeds ethics committee.

### Procedure

Participants sat at a desk with a touchscreen computer tablet (Lenovo ThinkPad Helix 2, 1920×1080 pixels, 11.6” screen, 60Hz refresh rate) placed directly in front of them and interacted with the screen using a stylus (sampled from the screen digitiser at a rate of 100Hz) in their chosen hand. Participants were shown a pictorial instruction sheet prior to starting the experiment that explained how to complete a single trial. Prior to starting the experiment, participants were instructed that each trial should be completed as quickly and accurately as possible. The core trial structure was the same across all three experiments and key differences for each study are detailed below and presented in Figure 1.

### Experiment 1: Biases in Sensorimotor Decision-making

In Experiment 1, we introduced a novel sensorimotor decision task in which participants were asked to select one of two possible routes around an obstacle to reach a target using a stylus on a tablet display. During each trial, participants had to stay within a 200mm high by 106mm wide workspace, displayed as a rectangle on the screen. Participants began each trial by placing the stylus on the screen and moving the cursor (5mm diameter circle) to a start-point (10mm diameter circle, horizontally central) at the bottom centre of the screen. Events in the scene (e.g. entering the start-point) were triggered when the perimeter of the cursor intersected the perimeter of another object. After a randomly sampled delay (generated by a random number generator) between 300-600ms (across all participants and experiments, there was a grand mean delay of 448ms (SD = 9ms), with the mean average delay across participants ranging from 423– 475ms), the start-point changed to a colour and shape combination, randomly selected from a list of three of each (cyan, magenta, yellow; circle, square, triangle); a check-point (10mm diameter circle, horizontally central) appeared in the top centre of the screen; and an obstacle (30mm wide x 10mm high rectangle) appeared equidistant between the start- and check-points, vertically central to the screen. The distance between the start and check-points was 140mm. Participants were instructed to remember the colour they were shown and move as quickly and accurately after stimulus display to the checkpoint. Participants were allowed up to 2 seconds ‘preparation time’ in the start-point. Upon leaving the start-point the coloured shape disappeared, and the participant had up to 500ms to reach the check-point.

Upon entering the check-point, the obstacle disappeared. The participant then had to wait in the check-point. After a randomly sampled delay (generated by a random number generator) between 300-600ms (across all participants and experiments, grand mean delay = 450ms, SD = 11ms, range = 418–476ms), three targets at the bottom of the screen appeared, spaced equally in the horizontal axis. Each target had an invisible 10mm diameter circle used to detect the cursor hitting. The vertical distance between the check-point and targets was 150mm, with 30mm spacing between targets horizontally. Each target had a randomly assigned colour and shape combination, selected by randomly shuffling the list of three colours and three shapes and allocating the combinations to each target. Participants were instructed to move as quickly and accurately to the target of the same colour that was shown at the start-point. Participants were allowed up to 2 seconds in the check-point. After leaving the check-point the participant had up to 1 second to reach the target.

A trial was successfully completed when the participant moved to the correct target. There were several ways to fail a trial: hitting the obstacle or task boundaries; spending too long in the start or checkpoint; moving too slowly between the start- and check-points, or moving too slowly between the check-point and target; leaving the checkpoint before the targets were revealed; and moving to the wrong coloured target. If a failure was triggered the trial was immediately terminated. Once a trial finished, visual feedback was presented for 1 second to indicate the outcome of a trial, where the target turning green indicated a successful trial, and an object turning red indicated a failed trial. For failed trials, the object that turned red indicated the type of failure (e.g. if the participant hit the obstacle, it would turn red). Across all trials a running score was shown in the top left of the screen, which increased by one after each successful trial. After visual feedback had been presented for 1 second, the start-point was shown on the screen, and participants were able to begin a new trial. During the experimental trials, the mean within-subject time between the start of successful trials was 4.17 seconds, and 3.67 seconds for unsuccessful trials.

The experiment comprised a total of 106 trials and took approximately 10 minutes to complete. This included 4 example trials, 6 practice trials, 9 baseline trials, and 87 experimental trials made up in 3 blocks of 29 trials, shown in Figure 1d. During the example trials, the instructor showed the participant a set of 4 standard trials including two successful trials and two failed trials. Participants were provided 6 practice trials to make sure they understood the task mechanics, which included text feedback after every trial to indicate the outcome, in addition to the regular visual feedback. A baseline block followed that aimed to make participants move as quickly as they could (while maintaining accuracy) by giving text feedback telling them they needed to move more quickly if their movement time between the start- and check-point was slower than their previous fastest time (following the baselining block, across all experiments the within-subject mean movement time between the start and check-point = 369ms, SD = 47ms, range = 249 – 452ms). During each of these three blocks, the obstacle was located horizontally central to the screen, and after the completion of each block, text was displayed for 10 seconds on the screen to indicate the participant was starting a new phase of the experiment.

The experimental trials were then organised into three blocks, presented to the participant as one uninterrupted block. The three conditions were where the obstacle’s horizontal position moved sequentially between trials from the left of the screen to the right (Rightward), from the right of the screen to the left (Leftward), and where the obstacle’s positions were randomly shuffled (Random). The order of the three conditions was randomly allocated per participant. Each obstacle position was presented once per block. Twenty-nine obstacle positions were used with extreme positions of −34.2mm and 34.2mm, with equally spaced jumps between each position.

The experimental task was developed using Unity (Unity Technologies, 2018; version 2018.1) and the Unity Experiment Framework (Brookes et al., 2019).

### Experiment 2: Decreasing Temporal Constraints

Experiment 2 was conducted to explore whether easing the temporal constraints placed on the decision-making system could attenuate hysteresis. In Experiment 2, participants were forced to wait while the stimuli were shown before executing the movement. Upon entering the start-point a red box appeared surrounding the start point and the participant’s cursor. While the red box was visible the participant was not allowed to leave the start-point or the trial would terminate in failure. In common with Experiment 1, there was a randomly sampled delay of between 300-600ms before the stimuli were presented. However, in Experiment 2 the red box remained on the screen after stimulus display. After 1.5 seconds the red box disappeared, and the participant completed the rest of the trial as described in Experiment 1. Differences between these Experiments are illustrated in Figure 1b. In Experiment 2, the mean within-subject time between the start of successful trials was 5.50 seconds, and 4.22 seconds for unsuccessful trials in the experimental block.

### Experiment 3: Reducing Task Cognitive Demands

Experiment 3 was conducted to explore whether reducing the cognitive demands associated with the task could attenuate hysteresis. Participants were presented with only one target after waiting in the check-point, with the colour shown at the start of the trial always matching that of the target. The remainder of the trial followed the same structure as Experiment 1 (differences illustrated in Figure 1c). In Experiment 3, the mean within-subject time between the start of successful trials was 3.96 seconds, and 3.30 seconds for unsuccessful trials in the experimental block.

### A Computational Model of Action Selection

Figure 2 shows a computational model where the selection of an action can be influenced by successful completion of a previous action (because the estimate of the probability of success is dynamically updated on a trial-by-trial basis).

The model weights the probability of action selection (going left or right around the obstacle) by the ‘value’ of each action, where we define value as a combination of the costs, rewards, and a bias term reflecting the increased probability of success from repeating the previous action. The model comprises four free parameters – a scaling parameter for each of the cost, bias, and reward terms, *s*_*c*_, *s*_*b*_, *s*_*r*_, respectively, which are used to bring the terms to the same scale so the relative importance of each can be compared, and the rate at which biases are accumulated, *r*, which has a value in the range [0, 1]. Formally, the ‘value’ of the action *i* is defined as:

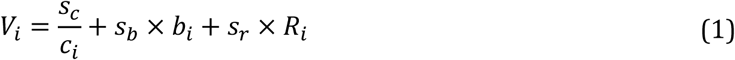

where *c* is the expected cost, *R* is the reward and *b* is the action bias accumulated over the previous trials. *R* is normalised about the minimum reward, so a total reward of 1 vs 3 becomes a normalised reward of 0 vs 2, so that the effect of an additional reward can be isolated. In this formulation, the expected cost of an action is evaluated perfectly but the model could be modified to include a distribution of the possible expected costs for an action. The reciprocal of the costs was used as a high cost should give a lower valuation of the action, whereas a high reward should give a higher valuation (thus, no transformation was used). The value was converted to selection probabilities using the soft argmax function. A random number was sampled from a uniform distribution between 0 to 1, and the left action selected if the random number was below the output probability and vice versa.

Action selection and outcome were used to update the biases associated with each action, according to the following reinforcement learning formula:

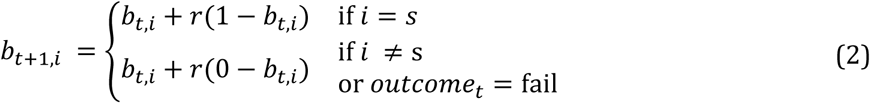

where *t* is the current trial number, *r* is the bias rate, and *S* is the selected action. It is assumed participants start with no bias and thus, biases are set to zero on the first trial.

In the experiments, the reward for successfully completing either action was the same, so the reward term in the model was not included. For simplicity, this model assumed the cost function was the path length of the movement trajectory. In fact, there is considerable debate (Todorov, 2004) within the sensorimotor research literature over the measure used for optimisation (candidates include path length, movement duration, normalised jerk, end-point variability, torque etc). We note that most of these factors co-vary and emphasise that there is a strong tendency for participants to select the shortest possible movement trajectory in unconstrained task settings (Tresilian, 2012). The observed paths were smooth and roughly symmetrical about the centre (see Figure S1 in Supplementary Materials) so path length was approximated by fitting the shortest parabola capable of connecting the start point to the target whilst passing the obstacle on either side. To aid model convergence, path lengths were divided by the minimum possible path length (140mm).

The model was fit to the choice data for the three experiments using Bayesian estimation via Stan (Carpenter et al., 2017; version 2.18.2). Each model was fitted with eight chains of 5,000 warmup samples and 5,000 iteration samples, giving 40,000 samples per posterior distribution. Convergence was assessed by visually inspecting chain behaviour and confirming the Gelman-Rubin statistic, 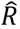, was below 1.1 (maximum 1.01) for all parameters (Gelman, 2004; Gelman & Rubin, 1992). Posterior distributions for each parameter were summarised using the 95% highest density interval (HDI), the 95% of most credible parameter estimates. The empirical priors used for model fitting are shown below. The scale of the model parameters was assessed by adjusting them until data containing hysteresis was observed. Aside from informing parameter scaling, the priors were then selected to be uninformative. Note that increasing the width of the priors doesn’t affect the results of the modelling.

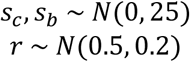

To check the model fit, datasets were simulated using each experimental list of obstacle positions from the real data. For each experiment, 10,000 samples of the posterior distribution were drawn, and each combination was used to simulate a new data set. For each combination of experiment and obstacle position, the probability of committing an error, extracted from the collected data, was used to simulate error trials. If a sampled number from a uniform distribution between 0 and 1 was below the error rate for the current experiment and obstacle position, the trial was classed as a failed trial and no action was selected. Each new dataset was summarised using a logistic fit for each of the three conditions, and at each obstacle position the minimum and maximum predicted probability of going right around the obstacle from all the logistic fits was taken, representing the credible range of possible data given the posterior distribution. The real data were then overlaid with the credible range to visualise the model fit.

### Data Analysis

The output from Unity included the experimental condition; the cursor position, sampled at 100Hz and output in millimetres; the timestamp when Unity’s physics engine detected participants had left the start-point; the position of the obstacle, output on the scale −1 to 1, where −1 indicated the obstacle touched the left wall and 1 indicated the obstacle touched the right wall; the direction participants moved around the obstacle, detected by Unity’s physics engine when participants moved past the leading edge of the obstacle; and the outcome of each trial. The extreme obstacle positions used were −0.9 and 0.9 (−34.2mm and 34.2mm respectively), which ensured participants could only go around the obstacle in one direction at these positions. The average error rate across the three experiments was 14.8%, 95% CI = [14.3, 15.4].

Stylus position data were filtered using a dual-pass Butterworth second-order filter with a cut-off frequency of 10Hz. To detect movement onset, the time where movement speed rose above 50mm/s closest to the Unity’s timestamp of the participant leaving the start-point was classed as movement start. For Experiment 1 and Experiment 3, reaction time (RT) was calculated as the difference in time between the obstacle being shown and movement start, whereas for Experiment 2 it was the difference between the red box disappearing and movement starting.

RT data were pre-processed by removing trials where no RT was present (117 trials, 0.7%), where RTs were lower than 100ms (36 trials, 0.2%, to account for participants anticipating stimuli presentation), then grouping within participant and condition and removing trials outside 2SD of the mean (826 trials, 4.9%), and then grouping by condition and experiment and removing participant’s conditions outside 2SD of the mean (240 trials, 1.4%). This RT data cleaning process was necessary to reduce heteroscedasticity and ensure normal residuals from models.

This process removed one participant (Experiment 1, mean RT = 608ms), six participant conditions (Experiment 1, mean RT = 587ms), and 1,219 trials in total (7.3% of observations) from the RT analysis. The remaining trials had a mean RT of 403ms. Of the trials removed, 7.2% were trials 1 and 2 in the experiment, likely because participants had only seen the obstacle presented central to the screen up to that point. Analysis performed on RT data was done on the inverse of RT to increase normality, and back-transformed when reporting. Choice data was pre-processed by removing trials where no movement past the obstacle was detected, removing 283 trials (1.7% of observations).

Analysis of the choice and RT data was performed using mixed-effect modelling, utilising the lme4 package in R (Bates et al., 2015; version 1.1-21). Following Barr et al. (2013), when the maximal random structure did not converge, the optimal random-effects structure was identified using forward model selection, with each mixed-effect model having a random intercept for participant. The effect of each variable was found using likelihood ratio testing, using the afex package (Singmann et al., 2019; version 0.23.0). Post-hoc comparisons were performed using the multcomp package (Hothorn et al., 2008; version 1.4.8), and corrections for multiple comparisons were made using the Bonferroni-Holm method. The MuMIn package (Barton, 2020; version 1.43.6) was used to report marginal *R*^2^ (variance explained by fixed effects), 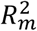, and conditional *R*^2^ (variance explained by fixed and random effects), 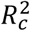, for the models (Nakagawa et al., 2017). The 95% confidence intervals for values are reported in square brackets throughout.

To examine changes in action selection, a mixed-effect logistic regression was performed. The fixed effects were the obstacle’s position, the condition, the experiment, and all combinations of the interactions between these variables. The model had a random intercept for each participant. While a model with a random slope for obstacle position converged, the single repetition of each condition led to an artificially steep main effect slope of obstacle position, so was not included in the model. The default condition was Random, the default experiment was Experiment 1, and the default obstacle position was 0. To understand whether hysteresis changed with experiment, the log-odds (LO) of going right around the obstacle at the central obstacle position was compared between conditions. Hysteresis was quantified as the increased LO of going right at the central obstacle position in the Rightwards condition compared to the Leftwards condition. We compared this across experiments.

To investigate changes in RT, a mixed-effect linear regression was performed. The fixed effects were the trial number in the block, the condition, the experiment, and all combinations of the interactions between these variables. The model had a random intercept for participant, with random slopes of condition. The default condition was Random and the default experiment was Experiment 1. The trial number in block was centred about the middle trial. To understand how RTs were affected, the estimated marginal mean (EMM) RT at the central obstacle position was compared between conditions. The difference in RT between the sequential conditions and Random was then compared between experiments.

The large sample size in Experiment 1 presented opportunity for more detailed analysis of hysteresis. We expected that, while there would be no global hysteresis in the Random condition, participants might exhibit biases within this block on a trial-by-trial basis. To explore this, a mixed effect logistic regression was performed to understand how much the previous trial biased the current trial inside the Random block. The fixed effects were the obstacle’s position, the prime condition – whether the participant went left (Left Previous) or right (Right Previous) on the previous trial, and the interaction between the two. The model had a random intercept for participant. As with the analysis of choice with condition, a random slope of obstacle position was omitted to avoid artificially inflating the main effect slope of obstacle position. The default prime condition was Left Previous. The default obstacle position was 0. To understand how selection was influenced by the previous trial, the LO of going right at the central obstacle position was compared between prime conditions.

As well as choices being biased by the previous trial, we found RTs were shorter when participants repeated their previous action when compared to switching action. To investigate the relationship between RT and hysteresis, a mixed-effect linear regression was performed on the Random condition, splitting the data by whether the participant switched or repeated the previous trial’s direction. The model had fixed effects of trial number, switch condition, and the interaction of the two, a random intercept for participant and a random slope of switch condition per participant. The default switch condition was repeated. The trial number in block was centred about the middle trial. The estimated mean RT at the central obstacle position was compared between switch conditions.

All statistical analyses and data processing were performed using custom-written scripts in R (R Core Team, 2018; version 3.5.2). Upon publication, all analyses code and model fits will be available through https://github.com/immersivecognition, and the complete dataset will be made available through the University of Leeds Data Repository.

## Results

### Choice Analysis

We first examined whether our group-level manipulations resulted in action selection biases. Hysteresis would result in participants going right around the obstacle more often in the Rightwards condition (where the obstacle moved from the left of the screen to the right between trials) and going left around the obstacle more often in the Leftwards condition (where the obstacle moved from the right of the screen to the left between trials), whereas the Random block (where the obstacle moved randomly between trials) should show no overall bias. We predicted that the degree of this bias would diminish when participants were provided with more planning time (Experiment 2) and when action execution was performed under a reduced cognitive task load (Experiment 3).

The experimental task was successful in revealing hysteresis (Figure 3a), with the effect was diminished in Experiment 2 (Figure 3b) and Experiment 3 (Figure 3c).

**Figure 3.**
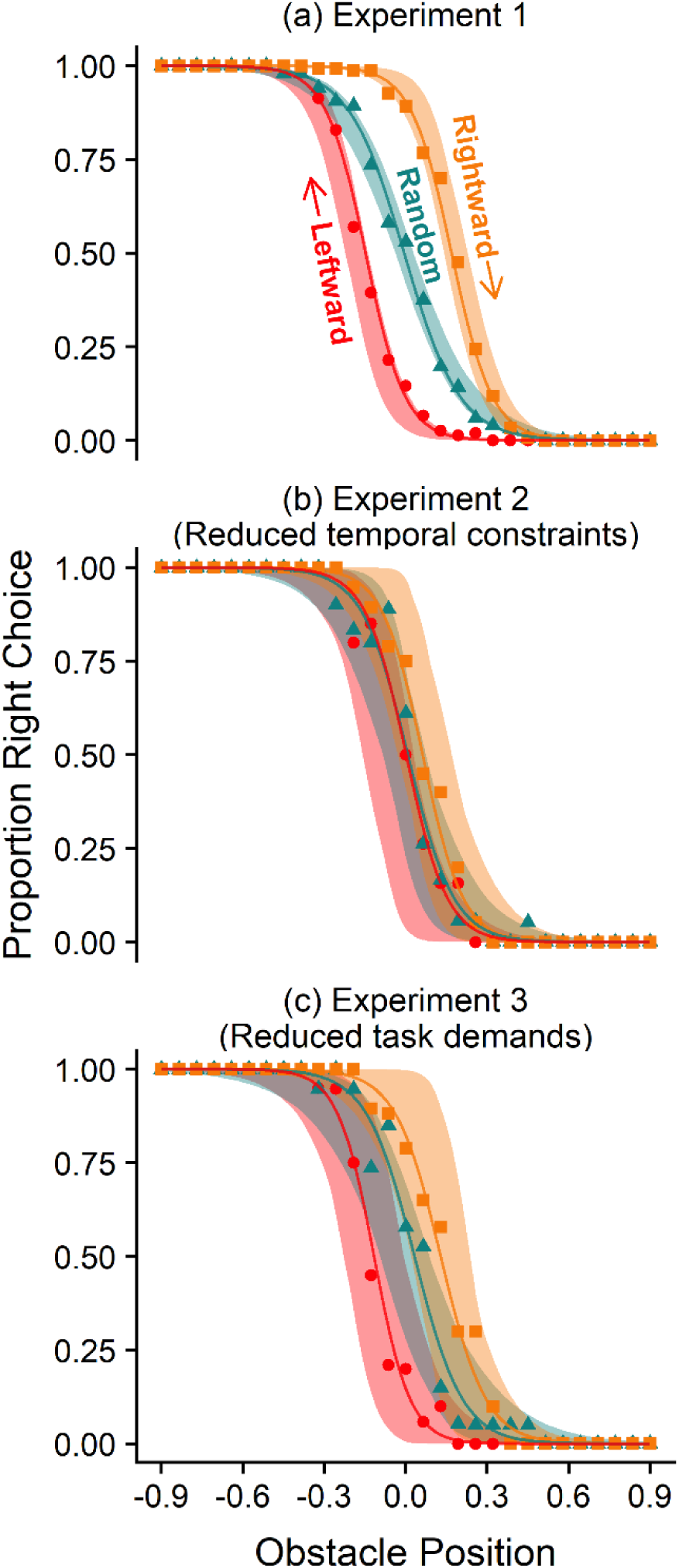
Comparison of experimental and simulated data for choices between experiments. The points and solid lines represent experimental data, and the ribbons represent simulated data. The points indicate mean proportion of participants who passed the obstacle on the right for each obstacle position. The solid lines represent the fit of a logistic regression for the experimental condition. Data was simulated using the decision-making model, with 10,000 samples of the posterior distribution used to simulate choices, and each new data set summarised with a logistic regression. The ribbon represents the minimum and maximum predicted probability of going right from the regressions of the simulated data. The conditions are Rightwards (where the obstacle moves from the left of the screen to right between trials), Random (where the obstacle moves randomly between trials), and Leftwards (where the obstacle moves from the right of the screen to left between trials.

We performed a mixed-effect logistic regression to predict the direction participants chose on a given trial. The model (χ^2^(17) = 18,206.60, p < 0.001, 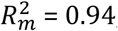, 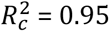) revealed a significant main effect of position (χ^2^(1) =4,627.52, p < 0.001) and condition (χ^2^(2) = 1,330.53, p < 0.001), but no significant effect of experiment (χ^2^(2) = 2.61, p = 0.272). There were significant interactions between position and condition (χ^2^(2) = 54.01, p < 0.001), and condition and experiment (χ^2^(4) = 94.53, p < 0.001), but no significant interaction between position and experiment (χ^2^(2) = 4.60, p = 0.100), or between position, condition and experiment (χ^2^(4) = 4.36, p = 0.359).

Bonferroni-Holm corrected comparisons were performed to see how the log-odds of passing the obstacle on the right changed with condition and experiment at the central obstacle position. In **Experiment 1**, participants were significantly more likely to go right in Rightwards compared to Random (LO = 2.52 [2.18, 2.86], p < 0.001), and significantly less likely to go right in Leftwards compared to Random (LO = −2.18 [−2.51, −1.85], p < 0.001), indicating participants were more likely to continue using the previous direction in the sequential conditions. Further, participants were more likely to go right in Rightwards compared to Leftwards (LO = 4.70 [4.28, 5.12], p < 0.001). This comparison gives the increased log odds of passing the obstacle on the right at the central obstacle positions between the sequential conditions, and is the measure of hysteresis used throughout.

In **Experiment 2**, where participants were forced to wait in the start-point for 1.5 seconds while the obstacle was shown before being allowed to move, they were significantly more likely to go right in Rightwards compared to Random (LO = 0.82 [0.05, 1.59], p = 0.026), but not in Leftwards compared to Random (LO = −0.07 [−0.80, 0.65], p = 0.811). Furthermore, participants were more likely to go right in the Rightwards trials compared to Leftwards (LO = 0.89 [0.11, 1.68], p = 0.023), indicating hysteresis was present, but to a lesser extent compared to Experiment 1.

In **Experiment 3**, where participants were presented with a reduced cognitive task load, they were significantly more likely to go right in Rightwards compared to Random (LO = 1.24 [0.44, 2.03], p < 0.001), and significantly less likely to go right in Leftwards compared to Random (LO = - 2.30 [−3.21, −1.39], p < 0.001). Participants were more also likely to go right in Rightwards compared to Leftwards (LO = 3.54 [2.54, 4.54], p < 0.001) indicating the presence of some hysteresis but attenuated relative to Experiment 1.

To investigate how the magnitude of hysteresis changed between the experiments, the increased log odds of going right in the Rightwards condition compared to the Leftwards condition were compared across experiments. Participants in Experiment 2 showed significantly less hysteresis than participants in Experiment 1 (LO = −3.81 [−4.70, −2.93], p < 0.001), and participants in Experiment 3 also showed significantly less hysteresis than participants in Experiment 1 (LO = - 1.16 [−0.09, −2.24], p = 0.012).

These results indicate that the experiment interventions designed to: (i) decrease temporal constraints and (ii) decrease cognitive task load reduced the magnitude of hysteresis in Experiments 2 and 3 respectively (relative to Experiment 1). We note that the impact seems to be more pronounced for planning time increase (Experiment 2) relative to task load reduction (Experiment 3).

Simulations from the action selection model were fit with the experimental data. The 95% HDI of the posterior distributions for each parameter are summarised in Table 1 (the full posterior distributions for each parameter are visualised in Figure S2 in Supplementary Materials). To understand whether the bias changed between experiments, the posterior distribution for the bias scaler parameter in Experiments 2 and 3 were subtracted from that of Experiment 1. This showed that the bias scaler estimate in Experiment 2 was lower than in Experiment 1 (mean difference = - 1.60, 95% HDI = [−1.18, −2.02]), and the estimate in Experiment 3 was also lower than in Experiment 1 (mean difference = −0.82, 95% HDI = [−0.38, −1.25]), indicating hysteresis was attenuated by reducing the contribution of biases, built up through previous successes and failures, towards action selection under reduced temporal constraints and cognitive load.

**Table 1.** Mean and 95% highest density interval (HDI) estimates of the decision-making model parameters.

| Experiment | Mean [95% HDI] |  |  |
| --- | --- | --- | --- |
|  | Cost scaler | Bias Scaler | Bias Rate |
| Experiment 1 | 39.26 [37.36, 41.24] | 2.69 [2.53, 2.86] | 0.32 [0.28, 0.36] |
| Experiment 2<br>(Reduced temporal constraints) | 41.96 [36.59, 47.84] | 1.09 [0.71, 1.48] | 0.43 [0.27, 0.62] |
| Experiment 3<br>(Reduced task demands) | 37.63 [32.94, 42.82] | 1.87 [1.48, 2.29] | 0.35 [0.21, 0.50] |

The posterior distributions were then used to simulate new data so that predictions from the model could be compared to the experimental data. Ten thousand samples of the posterior distribution were taken, and each used to simulate new responses to the experiments. Each new dataset was summarised with a logistic regression for each condition, and the upper and lower limit for these predictions used to visualise the model’s predictions. These predictions are shown as coloured ribbons in Figure 3 per experiment. Note that the observed selection probabilities lie within the range of the model’s predictions, with distinct separations between the two sequential conditions for Experiments 1 and 3 around the central obstacle positions, but for Experiment 2 the two sequential conditions share considerable overlap, consistent with the experimental data.

The results thus far indicate participants are biased towards repeating previously used action structures when the obstacle moves between trials in a sequential manner. While most hysteresis studies employ similar sequential trial designs, some have also found hysteresis when stimuli are varied randomly across trials (Jax & Rosenbaum, 2007; Valyear et al., 2018). Thus, we explored whether the participant’s selection on a current trial was biased by the direction they passed the obstacle on the previous trial in the Random block. Analysis of the data for repeated and switched trials from the Random block in Experiment 1 provided support for this idea (Figure 4).

**Figure 4.**
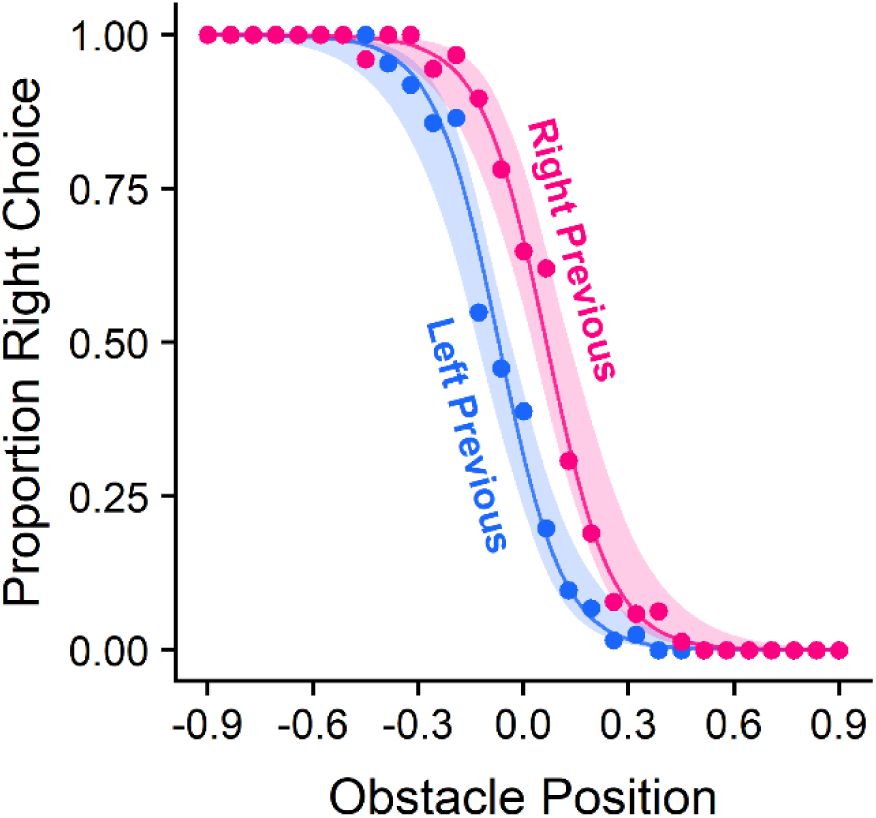
Empirical and simulated data for choices in Experiment 1 Random condition. The points and solid lines represent experimental data, and the ribbons represent simulated data. The points indicate mean proportion of participants who passed the obstacle on the left for each obstacle position. The solid lines represent the fit of a logistic regression for the experimental condition. Data were simulated using the decision-making model, with 10,000 samples of the posterior distribution used to simulate choices, and each new data set summarised with a logistic regression. The ribbon represents the minimum and maximum predicted probability of going right from the regressions of the simulated data. The conditions are Left Previous (where the participant passed the obstacle on the left on the previous trial), and Right Previous (where participants passed the obstacle on the right on the previous trial).

A mixed-effect logistic regression was performed on data from the Random block from Experiment 1 to predict the direction participants went on the current trial. The model (χ^2^(3) = 4,406.80, p < 0.001, 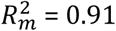, 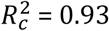) showed a significant main effect of position (χ^2^(1) = 2,215.09, p < 0.001) and prime condition (χ^2^(1) = 96.62, p < 0.001), but there was no significant interaction between position and prime condition (χ^2^(1) = 0.16, p = 0.688). A comparison was performed to see how the log-odds of passing the obstacle on the right changed with prime condition at the central obstacle position. Participants were significantly more likely to go right in the Right Previous condition compared to the Left Previous condition (LO = 1.51 [1.19, 1.83], p < 0.001). The increased log odds of going right were smaller for the two prime conditions when compared to the two sequential conditions from the analysis reported above - indicating biases accumulate over longer action sequences than just the previous trial.

To understand whether similar trial-to-trial biases emerged in the decision-making model, the parameter posterior distributions (10,000 samples) from Experiment 1 were used to simulate new datasets. The responses to the Random condition were then extracted, and each new dataset was summarised using a logistic regression per prime condition. The minimum and maximum predicted probability of going right from these regressions was used to visualise the model’s predictions and are represented as ribbons in Figure 4. Consistent with the observed experiment data, the model shows a distinct separation between the two prime conditions, with Right Previous cases being more likely to go right at the central obstacle positions.

### Reaction Times Analysis

Participants in Experiment 1 showed a reduction in RT in the sequential conditions compared to the Random condition (Figure 5a). Collapsing across all trials, the mean RT in the Random condition was 417ms [414, 419], compared to 387ms [385, 389] in the Rightwards condition and 388ms [386, 390] in the Leftwards condition. In Experiment 2, RTs were lower than in Experiment 1 and there seemed to be no RT benefits in the sequential conditions (Figure 5b), with a mean RT in the Random condition of 368ms [363, 373] compared to 381ms [375, 387] in the Rightwards condition and 380ms [374, 386] in the Leftwards condition. In Experiment 3, RTs were again lower than in Experiment 1 with a large RT benefit in the sequential conditions (Figure 5c). The mean RT in the Random condition was 379ms [374, 385] compared to 331ms [327, 336] in the Rightwards condition and 342ms [337, 347] in the Leftwards condition.

**Figure 5.**
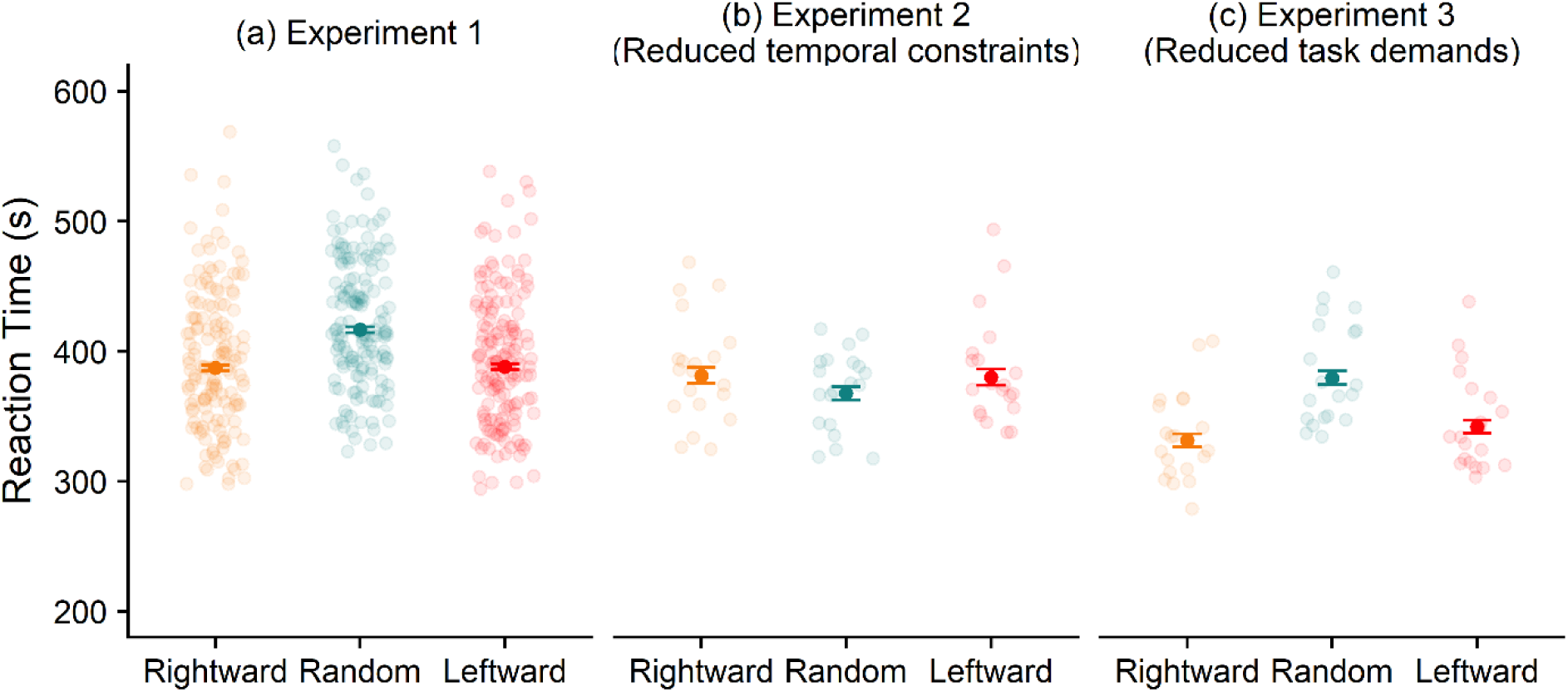
Comparison of RTs for each condition between experiments. The open circles show the mean RTs for each participant. The solid circles show the mean RT for each combination of condition and experiment across all participants, and the error bars show the 95% confidence intervals around the estimate of the mean. The conditions are Rightwards (where the obstacle moves from the left of the screen to the right between trials), Random (where the obstacle moves randomly between trials), and Leftwards (where the obstacle moves from the right of the screen to the left between trials).

To understand how RTs were affected by biases, a linear mixed-effect model was conducted to predict RTs on a given trial. The model [χ^2^(22) = 2,337.69, p < 0.001, 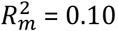, 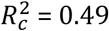] showed a significant main effect of trial (χ^2^(1) = 7.53, p = 0.006), condition (χ^2^(2) = 104.22, p < 0.001), and experiment (χ^2^(2) = 30.30, p < 0.001). There were significant interactions of trial and condition (χ^2^(2) = 28.45, p < 0.001), and condition and experiment (χ^2^(4) = 50.58, p < 0.001), but no significant interaction between trial and experiment (χ^2^(2) = 2.47, p = 0.291). There was a significant interaction between trial, condition and experiment (χ^2^(4) = 19.72, p < 0.001).

Bonferroni-Holm corrected comparisons were performed to see how RTs changed with condition and experiment at the middle trial in the block, where the benefits of hysteresis would be expected. In **Experiment 1**, the Random condition (EMM = 417ms [410, 425]) was significantly slower than the Leftwards (EMM = 389ms [381, 397], p < 0.001) and Rightwards conditions (EMM = 387ms [380, 396], p < 0.001), but there was no significant difference between the Leftwards and Rightwards conditions (p = 0.659). Participants were ∼30ms faster in the sequential conditions than in the Random condition, indicating RT savings from choice perseveration.

In **Experiment 2**, there were no significant differences between the Random (EMM = 368ms [352, 385]) and Leftwards condition (EMM = 380ms [361, 402], p = 0.151), the Random and Rightwards condition (EMM = 382ms [361, 404], p = 0.151), or between the Leftwards and Rightwards condition (p = 0.877).

In **Experiment 3**, the Random condition (EMM = 380ms [363, 398]) was significantly slower than the Leftwards (EMM = 342ms [326, 359, p < 0.001) and Rightwards conditions (EMM = 331ms [316, 348], p < 0.001), but there was no significant difference between the Leftwards and Rightwards conditions (p = 0.093). Participants were 40-50ms quicker in the sequential conditions than in the Random condition, showing savings that were marginally larger than those observed in Experiment 1.

To investigate whether the magnitude of RT reduction changed between experiments, we compared the difference in RT of the Random condition to the sequential conditions between experiments. The difference in RT between the Random and Leftwards conditions was significantly lower in Experiment 2 compared to Experiment 1 (p < 0.001), but significantly higher in Experiment 3 compared to Experiment 1 (p = 0.005). The difference in RT between the Random and Leftwards conditions was significantly lower in Experiment 2 compared to Experiment 1 (p < 0.001) but there was no difference between Experiment 3 and 1 (p = 0.074).

In the Random block of Experiment 1 participants showed a reduction in RT for repeated choices compared to switched choices (Figure 6). Collapsing across all trials and participants, the mean RT in the Repeated condition was 407ms [404, 410] compared to 430ms [427, 433] in the Switched condition.

**Figure 6.**
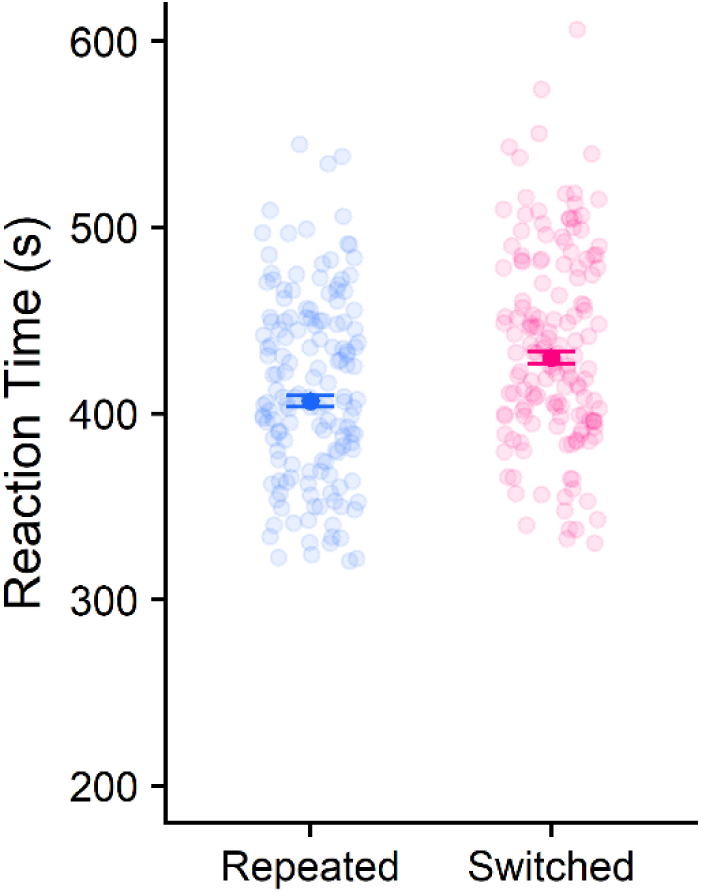
Comparison of RTs inside Experiment 1’s Random condition. The open circles show the mean RTs for each participant. The solid circles show the mean RT for each switch condition across all participants, and the error bars show the 95% confidence intervals around the estimate of the mean. The conditions are Repeated (where participants made the same choice on the current trial as on the last trial), and Switched (where participants switched choice from the previous trial).

To understand how RTs changed with switch condition on a given trial, a linear mixed-effect model was performed. The model [χ^2^(5) = 197.71, p < 0.001, 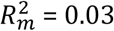, 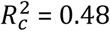] showed a significant main effect of trial (χ^2^(1) = 14.23, p < 0.001) and switch condition (χ^2^(1) = 97.79, p < 0.001), but no interaction between trial and switch condition (χ^2^(1) = 0.19, p = 0.663). A comparison was performed to see how RTs changed with switch condition at the middle trial in the block. The repeated condition (EMM = 407ms [400, 415]) was significantly faster than the switched condition (EMM = 429ms [421, 438], p < 0.001), indicating participants had an RT benefit from repeating only a single previous choice. The difference between the Switched and Repeated conditions was lower than the difference between the Random and the sequential conditions from the earlier analysis, indicating RT savings may be a cumulative process, building up across trials.

## Discussion

Our goal was to examine the bias shown by skilled adult humans towards selecting a previously selected action structure when alternative options that would be selected in *de novo* conditions were available. To this end, we created a simple obstacle avoidance aiming task. We found participants exhibited hysteresis effects when the obstacle moved systematically in one direction between trials. In blocks where the obstacle moved randomly, there was no global hysteresis effect. The random blocks did, however, show trial-by-trial biases - with action selection being influenced by the previous movement. We were also interested in exploring the impact on hysteresis of changing the temporal constraints of the task. The rationale for manipulating the temporal constraints was based on the growing evidence that human decision-making involves evidence accumulation processes. The existence of such processes suggests that humans may choose to act before a full evaluation of the costs associated with the available action options has been completed (i.e. as soon as an action reaches a threshold it is selected). It follows that providing a longer time period for decision-making might cause a different action to be selected (as the available time can be used to more fully evaluate the costs, reducing the reliance of the current decision on previous successes and failures). We used two manipulations to alter the temporal constraints of the task. In Experiment 2, we directly manipulated the temporal constraints by preventing action until a 1500ms time window had elapsed. In Experiment 3, we indirectly altered the constraints by decreasing the cognitive demands of the task (reasoning that less time spent identifying the task goal would provide more time for evaluating the respective costs of the available actions). The results showed that the hysteresis effect was practically eliminated in Experiment 2 and attenuated in Experiment 3.

Our investigation of hysteresis was motivated by our hypothesis that hysteresis is the naturally emergent property of a dynamical learning system that is operating in an uncertain world. In order to test our hypothesis we created a simple POMDP type computational model that incorporated dynamic probabilistic estimate updating. We used this model to simulate behavioural responses for the experimental tasks and found that it was able to capture the empirical data. This finding suggests that there is no need to invoke the existence of a bespoke ‘hysteresis function’ within the sensorimotor system, and provides support for our hypothesis that hysteresis is an emergent property of a dynamical learning system.

The results reported within this manuscript emphasise the dynamical nature of human sensorimotor decision making. The challenge for the human nervous system is to maintain optimal action selection in a noisy and uncertain world. The only way that the nervous system can maintain its efficiency is through an ongoing evaluation of the accuracy of its internal representation of the external world, and frequent updating of its probability estimates. The current findings suggest that this updating occurs on a trial-by-trial basis (though the biases we observed also accumulated across multiple trials). This paints a picture of a system that is continually adapting, and ensuring that its actions are precisely tailored to the external environment. This observation calls into question the classical distinctions between sensorimotor control and sensorimotor learning. It appears that human control systems appear stable because they have been refined over long periods of time through interactions with a world that obeys consistent rules described by Newtonian mechanics– but controllers are nevertheless updated continually on the basis of feedback from every interaction. Our findings also highlight the dynamics of decision-making in terms of the system needing to make choices under time constraints (where there are strong evolutionary pressures favouring species who react swiftly). In line with the existence of evidence accumulation processes, we found that the hysteresis effect was attenuated when the temporal constraints of the task were eased. This emphasises the tendency within the system to select the first action to cross a pre-specified threshold, favouring recently successful actions, rather than wait until the full costs of all options have been exhaustively evaluated. The success of such a strategy is witnessed by the fact that Homo sapiens remained standing after the evolutionary arms race of the past gigayear.

Our experiments have focussed on the sensorimotor system but the general phenomenon of hysteresis can be observed in other aspects of human behaviour. For example, hysteresis effects have been observed in perceptual decision-making tasks where choices are biased by previous decisions (Abrahamyan et al., 2016; Akaishi et al., 2014; Urai et al., 2019). The magnitude of the perceptual bias tends to depend on whether the previous decision was rewarded or not (Abrahamyan et al., 2016; Hermoso-Mendizabal et al., 2018). These hysteresis effects (typically described as choice-history biases) have been successfully implemented in evidence accumulation models of decision-making. We argue that the existence of hysteresis within the perceptual system can be explained through the same mechanisms that we used to account for hysteresis in sensorimotor decision-making (i.e. the presence of Bayesian type processes operating within the brain, where priors are continually updated with new sensory information to create posterior probability estimates). It is possible that similar mechanisms can account for reports of hysteresis in higher order cognition (often described as ‘perseveration’). There is a growing consensus that the sensorimotor system provides the phylogenetic and ontogenetic foundations of higher order cognition (Raw et al., 2019; Wilson, 2002). The postulated links between the sensorimotor and cognitive system might suggest a close relationship between sensorimotor hysteresis and perseveration type behaviours. This may prove a fruitful line of investigation for future studies.

Previous accounts of hysteresis have assumed that the sensorimotor system has a ‘hysteresis’ function whose purpose is to create an advantage when planning a new movement. It is argued that modifying a previously used plan would be more cognitively efficient than planning a new one from scratch, so hysteresis exists to improve planning efficiency (Meulenbroek et al., 1993; Rosenbaum et al., 2007; Schütz & Schack, 2019; Weiss & Wark, 2009), as indexed by reduced RTs when performing the same action as previously (Valyear et al., 2018). On the basis of previous reports we fully expected to find reduced RTs when the hysteresis effect was present. Moreover, we anticipated the presence of reduced RTs on the theoretical basis that evidence accumulation processes will cause actions to be selected more rapidly when there is a bias towards one action versus another. In line with these expectations, we observed a decrease in the average RT on the sequential trials in Experiment 1 relative to the random trials. In Experiment 2, participants were given a substantial time to select the goal directed action and the RTs were similar to the sequential trials in Experiment 1 regardless of trial type. In Experiment 3, the task demands were decreased and there was a commensurate reduction in RT (consistent with a large body of literature showing that RT is a function of task complexity). Notably, the sequential trials within Experiment 3 showed hysteresis (relative to the random trials within the experiment) and were associated with faster RTs than the random trials (producing the fastest RTs across all three experiments as predicted by the presence of hysteresis and the reduced task complexity).

The work presented within this manuscript addresses issues from the field of ‘Human-Like-Computing’ where researchers attempt to bridge the gap between models of human decision-making and the models used in artificial intelligence and robot motion control. Stochastic models of actions, observations, costs and rewards are the main tools used in modelling and planning robot motion, including tasks that involve reaching behind obstacles (Dogar & Srinivasa, 2012). An improved understanding of human decision-making can inform the development of such robot motion models. The identification of the hysteresis bias allows roboticists and computer scientists to decide whether their agents are operating within environments that are sufficiently constrained so that control schemes can seek to ameliorate hysteresis. Alternatively, hysteresis may suggest mechanisms through which a robotic agent can show human-like flexibility and adaptability in complex dynamic environments. Moreover, the identification of hysteresis as an emergent property can help improve the legibility and predictability available within human-robot interactions. It follows that investigations into human biases (such as hysteresis), and their formal description through mathematical models can be useful in robot motion and control. Thus, the approach adopted within this manuscript provides an interesting avenue for future investigations by roboticists, psychologists and computer scientists.

## Competing Interest

The authors declare that they have no competing financial or non-financial interests.

## Acknowledgements

We thank Peter Culmer and Ian Flatters for their contributions on preliminary work for this project. We also thank J. Ryan Morehead for comments on a recent version of the manuscript. Authors A.G.C., F.M and M.M-W hold Fellowships from the Alan Turing Institute. This project was funded by a grant from the EPSRC Human-Like Computing Call (EP/R031193/1).

## Notes

### Competing Interest Statement

The authors have declared no competing interest.

