## Supplemental Materials for "Getting stuck in a rut as an emergent feature of a dynamic decision-making system"

**Supplement A**

Participant's trajectory between the start- and check-points for all successful trials are shown in Figure S1.

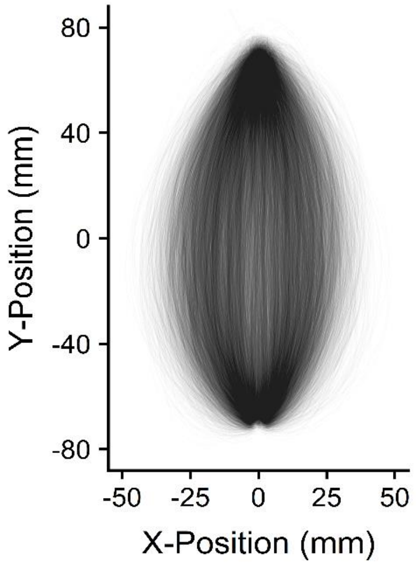


Figure S1. Trajectories of all trials from all participants between the start-point to check-point. Only trials where the participant reached the check-point are included.

**Supplement B**

The full posterior distribution for the model parameters is shown in Figure S2. The 95% highest density interval (HDI; the 95% of most credible parameter estimates) are shown underneath distribution. Notably, the cost scaler and bias rate are similar between all experiments, whereas the bias scaler reduces in Experiments 2 and 3, relative to Experiment 1.

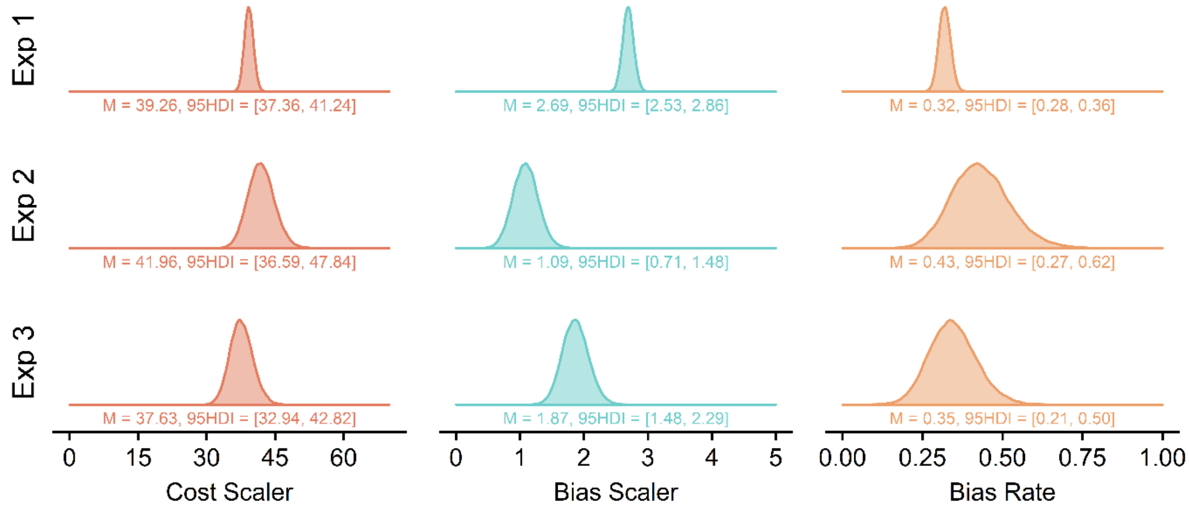


Figure S2. The full posterior distributions for the model parameters across the three experiments. The mean value and the 95% highest density interval (HDI) are shown underneath the density plot. Notably, the cost scaler and bias rate stays similar between experiments, but the bias scaler is reduced in Experiments 2 and 3 compared to Experiment 1, explaining the prediction of reduced hysteresis in the simulated data.
